# Potential involvement of protein phosphatase PP2CA on protein synthesis and cell cycle during SARS-CoV-2 infection. A meta analysis investigation

**DOI:** 10.1101/2023.06.02.543487

**Authors:** Luca P. Otvos, Giulia I.M. Garrito, Luciana E.S.F. Machado

## Abstract

Coronavirus disease 2019 is a multi-systemic syndrome that caused a pandemic. Proteomic studies demonstrate changes in protein expression and interaction involved in signaling pathways related to SARS-CoV-2 infections. Protein phosphatases are important for cell signaling regulation. Here we aimed to understand the involvement of protein phosphatases and the signaling pathways that may be involved during SARS-CoV-2 infection. Then, we carried out a metanalysis of protein phosphatase interaction directly or indirectly with viral proteins. Additionally, we analyzed the expression degree of protein phosphatases, and phosphorylation degree of intermediate proteins. Our analyses revealed that PP2CA and PTEN were the key protein involved in the cell cycle and apoptosis regulation, during SARS-CoV-2 infection. Showing it as potential target for COVID-19 control.

## Introduction

Coronavirus disease 2019 (COVID-19) is caused by infection from severe acute respiratory syndrome coronavirus 2 (SARS-CoV-2) that reached a pandemic level and was a massive global health threat. The pandemic Covid-19 has 676,609,955 global cases report, obtained on March 29^th^, 2023 at 10:45 am in the Coronavirus Resource Center of Johns Hopkins University (JHU) website (https://coronavirus.jhu.edu/map.html).

Covid-19, with its higher potential of viral transmission and high lethality worldwide, has no effective treatment, raising awareness of researchers to understand the disease mechanism, to develop potential drugs for disease treatment as well as to develop vaccine to prevent from infection. The output from this global effort resulted in about 200.000 scientific papers on “COVID-19” and “SARS-CoV-2” in nearly two years.

Although the Covid-19 had been described as a respiratory disease, it is better explained as a multisystem syndrome that affects different organs and tissues in the affected patients^1^. Multi-level assessment of transcripts, protein and post-translational modifications in cultered cells, human biopsies and fluids revealed a strong correlation between the degree of immune response activation and the disease severity^2–11^.

A common aspect revealed by these studies demonstrate that protein kinases are activated during cell signaling upon SARS-CoV-2 infection, suggesting that protein phosphorylation plays an important role to allow viral infection^6,11,12^.

Protein phosphatases and kinases are important to regulate cell signaling in the physiological and pathological functions, such as cell cycle control, cell survival, apoptosis, cancer, diabetes, cardiovascular diseases, and others. In these processes, kinases phosphorylate and phosphatases dephosphorylate their substrates, regulating the degree of protein phosphorylation ^13,14^ and controlling signaling duration (initiation, sustain and termination) and amplitude ^15–17^.

Protein phosphatases are classified in two families, according to their substrate specificities: protein serine threonine phosphatase (PPPs) and protein tyrosine phosphatases (PTPs). The PPPs is further divided in three classes: protein serine/threonine phosphatase (PPP), protein phosphatase metal-dependent (PPM) and Aspartate-based protein phosphatase (FCP/SCP). The PTPs family is further divided in three classes: classical protein tyrosine phosphatase, dual-specificity phosphatases (DUSPs) that are tyrosine, serine and threonine phosphatase, and Aspartate- or Histidine-based phosphatases. In addition, classical PTPs can be divided in two groups, cytoplasmic PTPs (cPTPs) and receptor PTPs (rPTPs).

Once, protein phosphatases regulate the cell signaling in different pathophysiological process, we hypothesized that protein phosphorylation is promoted during SARS-CoV-2 infection by down-regulating protein phosphatases, amplifying the signaling duration and maximum signaling amplitude leading to cell death. To accomplish this, we investigated here the protein phosphatase levels and interactions available in distinct public databanks.

## Material and methods

### In silico analysis

Here we provide an in silico metanalysis that takes profit of the reanalysis of 26 scientific papers and their respective databases^2–7,9,11,12,18–34^ (**Figure 1, Supplementary Table 1**). Specifically, we selected scientific papers containing proteomic analyses performed in different model cells or using blood cells, plasma, urine and lung tissues from SARS-CoV-2 infected patients. The search for the expression, and/or interaction information of protein phosphatases and phosphorylation level of intermediate proteins (**Supplementary Table 2-4**) was obtained from both the supplementary files made available by the authors and in PRIDE and ProteomeXchange databases ^35^.

**Figure 1:**
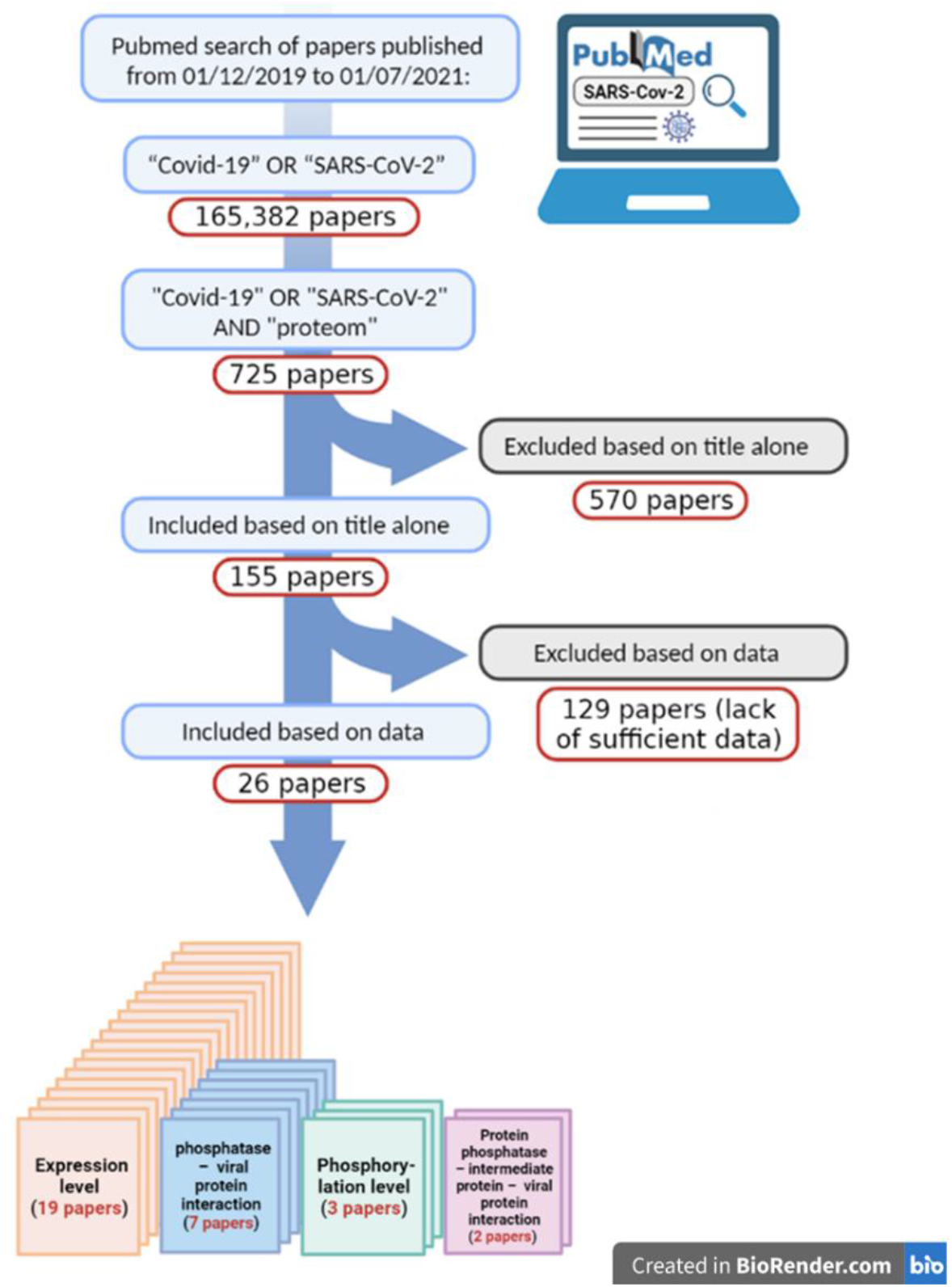
Flowchart of paper selection based on defined inclusion and exclusion criteria. The scientific papers were searched in the Pubmed using the words “Covid-19” OR “SARS-CoV-2”, resulting in 165,382 papers and refined adding AND “proteom”, resulting in 725 papers from 01/12/2019 to 01/07/2021. Based on the abstract, 570 papers were excluded, resulting in 155 papers to be analyzed. Due the lack of sufficient data, as the fold change and statistic analysis, 129 papers were excluded. Finally, 26 papers were analyzed, being 19 papers for expression level of protein phosphatases, 7 papers for protein phosphatase-viral protein interaction, 3 papers for phosphorylation level of intermediate proteins and 2 papers for protein phosphatase – intermediate protein – viral protein interaction.

### Protein-protein interaction network

The generation of interaction networks between host and viral proteins was carried out by the Cyoscape software using data extracted from publicly available databases. (**Supplementary Table 3**) ^36^.The function of proteins interacting with protein phosphatases and viral proteins were analyzed using the Uniprot program ^37^. We povide a preliminary dataset containing all proteins predicted to interact with protein phosphatases and viral proteins. This dataset was used as an input into the Cytoscape program using MCode and ClueGo plugin ^38,39^ in order to identify the signaling pathways they are involved.

### Statistical analysis

The statistical analysis and the graphs obtained from the expression and the phosphorylation levels of protein phosphatases and intermediate proteins were obtained using the GraphPad Prism 9 program. We used the Fold Change data and log_10_ p-value values for each sample extracted. These data refer to the ratio of samples from control groups, and samples from infected groups, accordingly with the equation FC = log_2_(infected/control), where FC ≥ 1.0 represent the higher level of protein phosphorylation compared to control. FC ≤ 1.0 represent the lower level of protein phosphorylation compared to control (**Supplementary Table 2, 4**). The data with statistical difference were considered when p < 0.05 (log_10_ pvalue > 1.3).

## Results and Discussion

### Expression levels of protein phosphatases during SARS-CoV-2 infection

To understand the role of host protein phosphatases during SARS-CoV-2 infection, we analyzed the expression level of protein phosphatases in different datasets. Those studies were carried out in cell models or samples from confirmed SARS-CoV-2 infected patients (urine, blood cells, plasma, pulmonary tissues) ^2–5,7,9,11,12,19,21,26–34^. Interestingly, we observed a similar profile of protein phosphatase expression level among their sub-families between infected and control sample. This was independent of the analyzed model (model cell, patient tissue or cell and patient fluids and secretions), which consist of higher expression for PTPs and lower expression for PPPs.

In details, for the cell models, mostly of the expression level differences between infected and control cells were observed in the liver tumor cell (Huh7), which present higher expression level for cPTPs (numbers 15, 20, 24 and 27 in **Figure 2A**), specially for rPTPs (numbers 14, 16-19, 26, 29, 31, 32 in **Figure 2A**) and lower expression level for PPM (numbers 1, 2, 5, 7, 8 in **Figure 2A**). While the other HEK293 kidney derived human cell line, A459 lung derived human cell line and colorectal adenocarcinoma Caco-2 cells, showed higher expression level only in the PPP1CB (number 28 in **Figure 2A**) and PPP4C (number 22 in **Figure 2A**) for HEK293 cells, PTPN2 (number 25 in **Figure 2A**) for A459 cells. And lower expression level only in the PTPN13 (number 10 in **Figure 2A**) and PTPRF (number 9 in **Figure 2A**) for A459 cells, and PTPRF for Caco-2 cells (**Figure 2A, Supplementary Table 2**).

**Figure 2:**
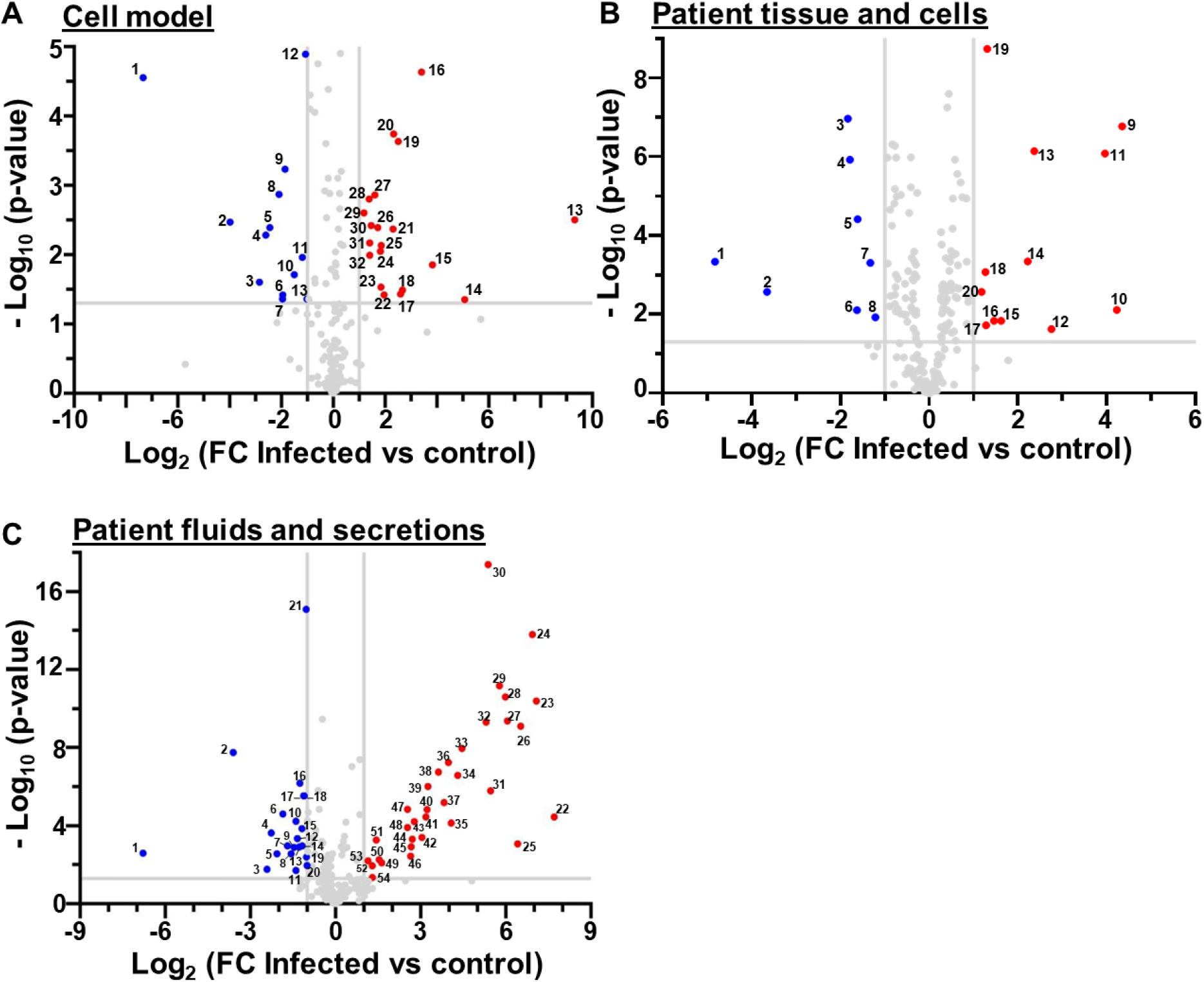
Volcano plot of expression levels of protein phosphatases. FC means the fold change of expression level of the ratio between infected cells by SARS-CoV-2 versus no-infected (control) in (A) cell model, (B) patient tissue and cells, (C) patient fluids and secretions. FC ≥ 1.0 represent the higher level of protein expression compared to control (red). FC ≤ 1.0 represent the lower level of protein expression compared to control (blue). FC = log2(infected/control). Graph was plotted using GraphPadPrism 9.0. Legend numbers in A, lower level of protein phosphatase expression: 1 - PDP2 (Huh7 cells); 2 - PPP1CC (Huh7 cells); 3 - PTPRU (Huh7 cells); 4 - DUSP6 (Huh7 cells); 5 - PPM1E (Huh7 cells); 6 - DUSP22 (Huh7 cells); 7 - PPM1L (Huh7 cells); 8 - PPM1H (Huh7 cells); 9 - PTPRF (A459 cells); 10 - PTPN13 (A459 cells); 11 - PTPN1 (Huh7 cells); 12 - PTPRF (Caco-2 cells); 13 - PTPMT1 (Huh7 cells) and higher level of protein phosphatase expression: 13 - PPM1J (Huh7 cells); 14 - PTPRS (Huh7 cells); 15 - PTPN21 (Huh7 cells); 16 - PTPRF (Huh7 cells); 17 - PTPRZ1 (Huh7 cells); 18 - PTPRH (Huh7 cells); 19 - PTPRJ (Huh7 cells); 20 - PTPN12 (Huh7 cells); 21 - PDP1 (Huh7 cells); 22 - PPP4C (HEK293 cells); 23 - DUSP16 (Huh7 cells); 24 - PTPN3 (Huh7 cells); 25 - PTPN2 (A459 cells); 26 - PTPRM (Huh7 cells); 27 - PTPN6 (Huh7 cells); 28 - PPP1CB (Huh7 cells); 29 - PTPRK (Huh7 cells); 30 - PPP1CB (HEK293 cells); 31 - PTPRG (Huh7 cells); 32 - PTPRA (Huh7 cells). Legend numbers in B, lower level of protein phosphatase expression: 1 - PTPRC (Lung); 2 - PTPN6 (Lung); 3 - PPM1M (PBMC cells); 4 PPM1B (PBMC cells); 5 - PPM1K (PBMC cells); 6 - PPP2CA (Lung); 7 - PGAM1 (Spleen); 8 - PGAM1 (Lung) and higher level of protein phosphatase expression: 9 - PPM1E (PBMC cells); 10 - DUSP8 (PBMC cells); 11 - PGAM4 (PBMC cells); 12 - PPM1B (Lung); 13 - PTPRN (PBMC cells); 14 - PPM1H (PBMC cells); 15 - PTPRU (PBMC cells); 16 – PTPRH (PBMC cells); 17 - PTPN11 (Lung); 18 - PTPN3 (PBMC cells); 19 - PTPN1 (Renal cortex); 20 - PTPRZ1 (Lung). Legend numbers in C, lower level of protein phosphatase expression: 1 - PTPRN (Urine); 2 - PTPMT1 (Bronchoalveolar lavage); 3 - PTPRO (Bronchoalveolar lavage); 4 - PTPRJ (Plasma); 5 - PTPRJ (Plasma); 6 - PTPN22 (Bronchoalveolar lavage); 7 - PPP1CC (Bronchoalveolar lavage); 8 - PTPDC1 (Bronchoalveolar lavage); 9 - PTPN9 (Bronchoalveolar lavage); 10 - PTPN18 (Bronchoalveolar lavage); 11 - PTPRM (Bronchoalveolar lavage); 12 - PTPN6 (Bronchoalveolar lavage); 13 - DUSP12 (Bronchoalveolar lavage); 14 - PPP1CA (Bronchoalveolar lavage); 15 - PPP3CB (Bronchoalveolar lavage); 16 - PPP6C (Bronchoalveolar lavage); 17 - PTPA (Bronchoalveolar lavage); 18 - PTPRA (Bronchoalveolar lavage); 19 - DUSP3 (Bronchoalveolar lavage); 20 - PTPRJ (Plasma); 21 - PTPRF (Plasma) and higher level of protein phosphatase expression: 22 - DUSP13 (Bronchoalveolar lavage); 23 - PTPRZ1 (Bronchoalveolar lavage); 24 – PTPRG (Bronchoalveolar lavage); 25 - PTPRQ (Bronchoalveolar lavage); 26 - PTPRT (Bronchoalveolar lavage); 27 - PTPRF (Bronchoalveolar lavage); 28 - PTPRK (Bronchoalveolar lavage); 29 - PTPN14 (Bronchoalveolar lavage); 30 - PTP4A1 (Bronchoalveolar lavage); 31 - PTPRD (Bronchoalveolar lavage) 32 - DUSP2 (Bronchoalveolar lavage); 33 - PTPRS (Bronchoalveolar lavage); 34 - PTPRN2 (Bronchoalveolar lavage); 35 - PTPRH (Bronchoalveolar lavage); 36 - DUSP14 (Bronchoalveolar lavage); 37 - PTPRU (Bronchoalveolar lavage); 38 - DUSP4 (Bronchoalveolar lavage); 39 - DUSP1 (Bronchoalveolar lavage); 40 - DUSP5 (Bronchoalveolar lavage); 41 - PTP4A3 (Bronchoalveolar lavage); 42 - DUSP16 (Bronchoalveolar lavage); 43 - PTPN13 (Bronchoalveolar lavage); 44 - PTPN21 (Bronchoalveolar lavage); 45 - DUSP19 (Bronchoalveolar lavage); 46 - PTPN3 (Bronchoalveolar lavage); 47 - PTPRE (Bronchoalveolar lavage); 48 - PGAM4 (Bronchoalveolar lavage); 49 - PGAM1 (Blood and Urine); 50 - PTPN7 (Bronchoalveolar lavage); 51 - PTPN18 (Nasopharinx swab); 52 - PTEN (Bronchoalveolar lavage); 53 - PTPRJ (Bronchoalveolar lavage); 54 - PGAM1 (Blood and Urine).

In the patient samples (fluids/cells), the protein phosphatase with higher expression level were for PTPs (numbers 10, 13, 15, 16, 18 in **Figure 2B**) for PBMC cells, (numbers 17, 20 in **Figure 2B**) for lung tissue, PTPN1 (number 19 in **Figure 2B**) for renal cortex. While the protein phosphatase with lower expression level were PPM1B (number 4 in **Figure 2B**), PPM1K (number 5 in **Figure 2B**) and PPPM1M (number 3 in **Figure 2B**) for PBMC cells, PGAM1 (number 8 in **Figure 2B**), PPP2CA (number 6 in **Figure 2B**), PTPN6 (number 2 in **Figure 2B**) and PTPRC (number 1 in **Figure 2B**) for lung tissue and PGAM1 (number 7 in **Figure 2B**) for spleen (**Figure 2B, Supplementary Table 2**).

In the patient fluids and secretion, the protein phosphatase with higher expression level were cPTPs (numbers 29, 30, 41, 43, 44, 46, 50 in **Figure 2C**), rPTPs (numbers 23-28, 30, 31, 33-35, 37, 47, 53 in **Figure 2C**) and DUSPs (numbers 22, 32, 36, 38-40, 42, 45, 52 in **Figure 2C**) for bronchoalveolar lavage fluid, PTPN18 (number 51 in **Figure 2C**) for nasopharynx swab, PGAM1 (number 54 in **Figure 2C**) for blood and urine. While the protein phosphatase with lower expression level were PPPs (number 7, 14-16 in **Figure 2C**), PTPs (numbers 2, 3, 6, 8-13, 17, 18 in **Figure 2C**) for bronchoalveolar lavage fluid, PTPRF (number 21 in **Figure 2C**) and PTPRJ (number 20 in **Figure 2C**) for plasma, PTPRN (number 1 in **Figure 2C**) for urine (**Figure 2C, Supplementary Table 2**).

In summary, we observed that the model cell Huh7 and the bronchoalveolar lavage presented the protein tyrosine phosphatase family (DUSPs, non-receptor and receptor) as PTPN12 and PTEN highly expressed, while the protein serine threonine phosphatase canonical, as PPP1CA, PPP1CC, PPP2CA and metal-dependent were lower expressed in the same samples as well as in the lung tissue.

### Protein phosphatases interact directly with viral proteins

Given the importance of protein phosphatases to host cell, we analyzed the interaction network of these proteins with viral proteins, as well as the interaction with other proteins that interact with protein phosphatases and viral proteins (hereafter referred to as intermediate proteins), during SARS-CoV-2 infection. To accomplish this, we searched scientific papers with protein interaction data during infection, and identified the presence of protein phosphatases that interact directly with the virus, or with other intermediate proteins that interact with viral proteins^4,11,18,20–25^.

SARS-CoV-2 virus belongs to linage of β-COVs, formed by a single stranded positive sense RNA molecule surrounded by an envelope. The genome of SARS-CoV-2 is composed by a 5’capped mRNA, poly-A tail at 3’end, 5’and 3’ UTR region (untranslated region), code gene for accessory proteins; and the code gene to four structural proteins, including spike (S), membrane (M), glycoprotein envelope (E) and nucleocapsid (N) proteins; and 11 open reading frames (ORFs), which are orf1a and orf1b that containing the code genes for sixteen non-structural proteins (nsp1-16), and orf3a, orf3b, orf6, orf7a, orf7b, orf9b, orf9c and orf10;

The following SARS-CoV-2 structural proteins are important for viral assembling and budding: M protein is composed of three transmembrane domain and define the shape of the viral envelope; E protein is a transmembrane protein with ion channel activity; S protein is a trimeric glycoprotein that interacts with host cell receptors to allow the viral entry; and N protein wrap the RNA genome. On the other hand, the non-structural proteins are important for viral transcription and/or replication (nsp7, nsp8, nsp9, nsp10, nsp13, nsp14, nsp15, nsp16, nsp11 and nsp12 that is an RNA-dependent RNA polymerase), blocking the host innate immune response (nsp1, nsp3), cleaving of viral polyprotein (nsp5), acting as scaffold protein (nsp4, nsp6), or having unknow function (nsp2) ^40,41^. At last, the accessory proteins are important to help the viral replication, virus release and modulate the host immune response.

To identify the host protein target for SARS-CoV-2 proteins, expression of 26 viral proteins in HEK-293T human cells identified 332 high confidence human protein interactions with SARS-CoV-2 proteins, by affinity purification and mass spectrometry ^18^. By searching in this data bank, we found thirteen protein phosphatases interacting directly with viral proteins, being six protein serine/threonine phosphatases, PPP1CA, PPP1CB, PPP1CC, PPP3CA, PPP6C and PGAM5; and seven are protein tyrosine phosphatases (PTP), involving two cPTP (PTPN1 and PTPN11), two rPTP (PTPRK and PTPRF), and three DUSP (DUSP11, DUSP14 and DUSP23/PTPMT1). Mostly of those identified PPPs can bind to several viral proteins, while PTPs seems to be more specific, binding to few viral proteins (**Figure 3A, Supplementary Table 3**).

**Figure 3:**
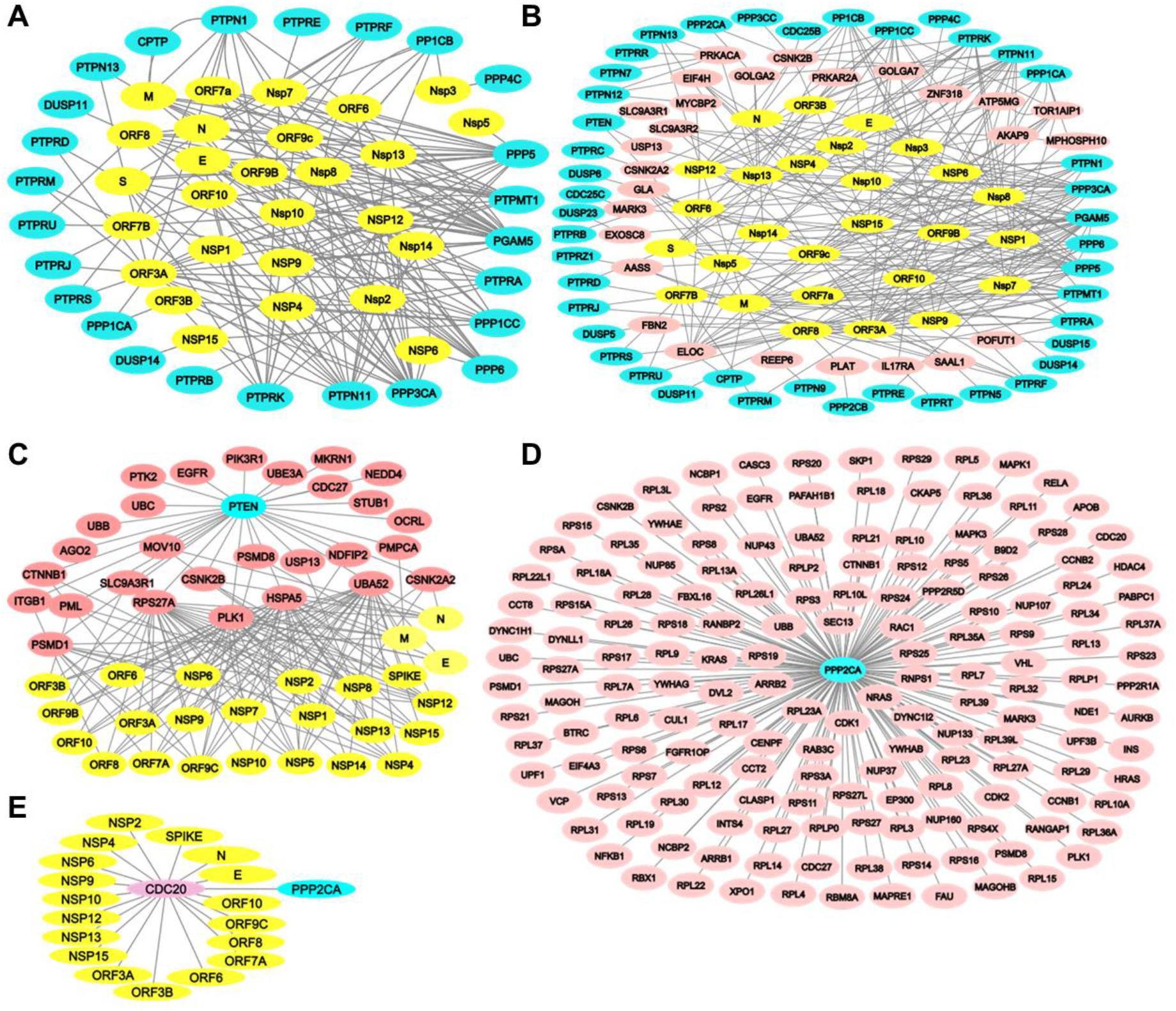
Protein-protein interaction network. (A) PPI between protein phosphatases (cyan) and SARS-CoV-2 viral proteins (yellow). (B) PPI between protein phosphatases (cyan) and intermediate proteins (magenta), which interact with SARS-CoV-2 viral proteins (yellow). (C) PPI of PTEN (cyan) interacting with intermediate proteins (magenta), which interact with SARS-CoV-2 viral proteins (yellow). (D) PPI of PP2CA (cyan) interacting with intermediate proteins (magenta), which interact with SARS-CoV-2 viral proteins (yellow). (E) Example of a intermediate protein (magenta) interacting with SARS-CoV-2 viral proteins (yellow) and PP2CA (cyan).

Using A459 lung carcinoma cells expressing individual SARS-CoV-2 and analyzed by affinity purification and mass spectrometry analysis, we identified interaction of seven protein tyrosine phosphatases PTPMT1, PTPN11, PTPRA, PTPRF, PTPRJ, PTPRM, PTPRS with ORF3 and/or ORF7B of viral proteins ^11^ (**Figure 3A, Supplementary Table 3**). We then mined a curated dataset of host and viral proteins interaction obtained from 151 publications. This dataset was obtained by International Molecular Exchange (IMEx) Consortium curators and returned 4400 interactions for those proteins described above. From these interactions, only PPP1CB was identified to interact with SARS-CoV-2 proteins (nsp8, nsp10, nsp13 and orf9b) ^25^ (**Figure 3A, Supplementary Table 3**). Additionally, Li et al (2021) overexpressed the SARS-CoV-2 gene in HEK293 cell and identified 286 cellular proteins interaction using affinity purification and mass spectrometry. From this dataset, we found PTPRD interacting with the viral protein orf3a of SARS-CoV-2 ^3^. While the analysis usign proximity proteomics that identified 2422 proteins, we identified two PTP (PTPN1 and PTPN13) and two PPP (PPP1CC, PPP3CA) interacting with some viral proteins ^24^ (**Figure 3A, Supplementary Table 3**). Expression of viral proteins and subsequent immunoprecipitation and mass spectrometry in human bronchial epithelial cells (16HBEo^-^) revealed that 189 host proteins interact with viral proteins. From these data, we identified eleven protein phosphatases, being two PPPs (PPP5, PPP6), eight PTPs (PTPRA, PTPRB, PTPRD, PTPRE, PTPRF, PTPRU PTPMT1, CPTP) and the histidine-based protein phosphatase phosphoglucomutase 5 (PGAM5) interacting with viral proteins ^23^ (**Figure 3A, Supplementary Table 3**).

The above mentioned protein phosphatases interacting with viral protein were grouped in five functional clusters: a) serine/threonine phosphatases activity, b) protein tyrosine phosphatase activity, c) transmembrane receptor protein phosphatase activity, d) DUSPs activity, and e) focal adhesion assembly. Furthermore, the proteins networks evidenced in these works were mapped into important cellular pathways affected by each viral proteins in the HEK293^3^: i)ATP biosynthesis and metabolic processes correlate with M viral protein; ii) mRNA transport with N viral protein; iii) melanoma differentiation-associated protein 5 and retinoic acid-inducible gene I RNA sensing signaling with Nsp1 viral protein; iv) nucleotide-excision repair with Nsp4 viral protein; v) protein methylation and alkylation with Nsp5; vi) translation initiation with Nsp9; vii) cellular amino acid metabolic process and neutrophil chemotaxis with Nsp10; viii) reactive oxygen species metabolic process with Nsp14; ix) Golgi to plasma membrane transport with Nsp16; x) endoplasmic reticulum stress and IFN or IL-6 signaling pathways with Orf3a; xi) mRNA transport and NF-κB pathways with Orf6; or in the A549 cells^11^: xii) mitochondrial dysregulation by ORF9b; xiii) innate immunity by ORF7b; xiv) stress response components and DNA damage response mediators by ORF7a; and xv) cholesterol metabolism by NSP6.

### Protein phosphatases interact with intermediate proteins

Although some studies show the direct interaction between protein phosphatases and viral proteins, other studies present protein-protein interaction (PPI) information describing intermediate proteins that made this binding. From that, we investigate the interactions between protein phosphatases and intermediate proteins, that interact directly with viral proteins. These interactions generate a network between many proteins that are important in a specific cell signaling pathways during SARS-CoV-2 infection once protein-protein interactions (PPIs) drive the cellular signal transduction mechanisms.

A protein-protein interaction prediction (HiPPIP) was implemented to assembly the interactome of host and viral proteins, generating a set of novel PPIs through a string-extended PPI network, which should be important during SARS-CoV-2 infection and replication ^20^. These data were extracted from 332 human proteins that interacted with the SARS-CoV-2 proteins analyzed by Gordon et al., 2020 and was obtained 4408 host proteins that were probably involved in 6076 interactions with viral proteins^20^.

Analyzing the interactome list, we identified twenty-seven protein phosphatases and twenty six intermediate proteins that interact with viral proteins. The protein phosphatases identified, six are PPPs (PPP2CA, PPP1CB, PPP1CC, PPP2CA, PPP2CB, PPP3CC) and twenty-one PTPs. Of these, eight cytosolic PTPs (PTPN1, PTPN5, PTPTN7, PTPN9, PTPN11, PTPN12, PTPN13, CPTP), six receptor PTPs (PTPRA, PTPRC, PTPRK, PTPRR, PTPRT, PTPRZ1) and seven DUSPs (DUSP5, DUSP6, DUSP15, DUSP23, PTEN, CDC25B and CDC25C) (**Figure 3B, Supplementary Table 3**). As a result, twenty-one new protein phosphatases interacting with intermediated proteins were identified in the interactome of SARS-CoV-2 infection.

Nadeau et al (2020) analyzed the viral-host PPI network obtained from Gordon et al., 2020 and performed an in-depth computational analysis of the interactome of SARS-CoV-2 using the GoNet tool. This tool evaluated the clustering of GO terms in PPI networks to identify biological processes and protein complexes, which involve three layers of interaction: *i)* viral-host PPI network, SARS-CoV-2 proteins interacting with human proteins; *ii)* string-augmented PPI network, interactions between human proteins in the viral-host PPI network; *iii)* string-extended PPI network, proteins that bind with human interactors of SARS-CoV-2 proteins ^22^. Analyzing these dataset, the protein serine/threonine phosphatase 2A catalytic subunit alpha (PPP2CA) was identified to bind with 166 human interactors of SARS-CoV-2 proteins and PTEN interacting with 28 human interactors of SARS-CoV-2 proteins (**Figure 3C-E, Supplementary Table 3**).

To analyze the biological functions of protein phosphatases interacting with intermediate proteins, we used MCode and ClueGo plugins in the Cytoscape software. The Mcode plugin identified seven function clusters, being three of them predominant (**Figure 4**). It is important to highlight that these software create networks based in nodes of interaction, being the protein with highest interaction observed as the central node and providing a higher percentage related to its function^38^. In the first cluster, most proteins were shown to be involved in four signaling pathway: *i)* SRP-dependent cotranslational protein targeting to membrane, *ii)* modulation by symbiont of host defense response, *iii)* ubiquitin ligase inhibitor activity and *iv)* ribosome assembly (**Figure 4A,D**). Three of these pathways were related to protein synthesis, while the other is related to the immunity defense against the virus.

**Figure 4:**
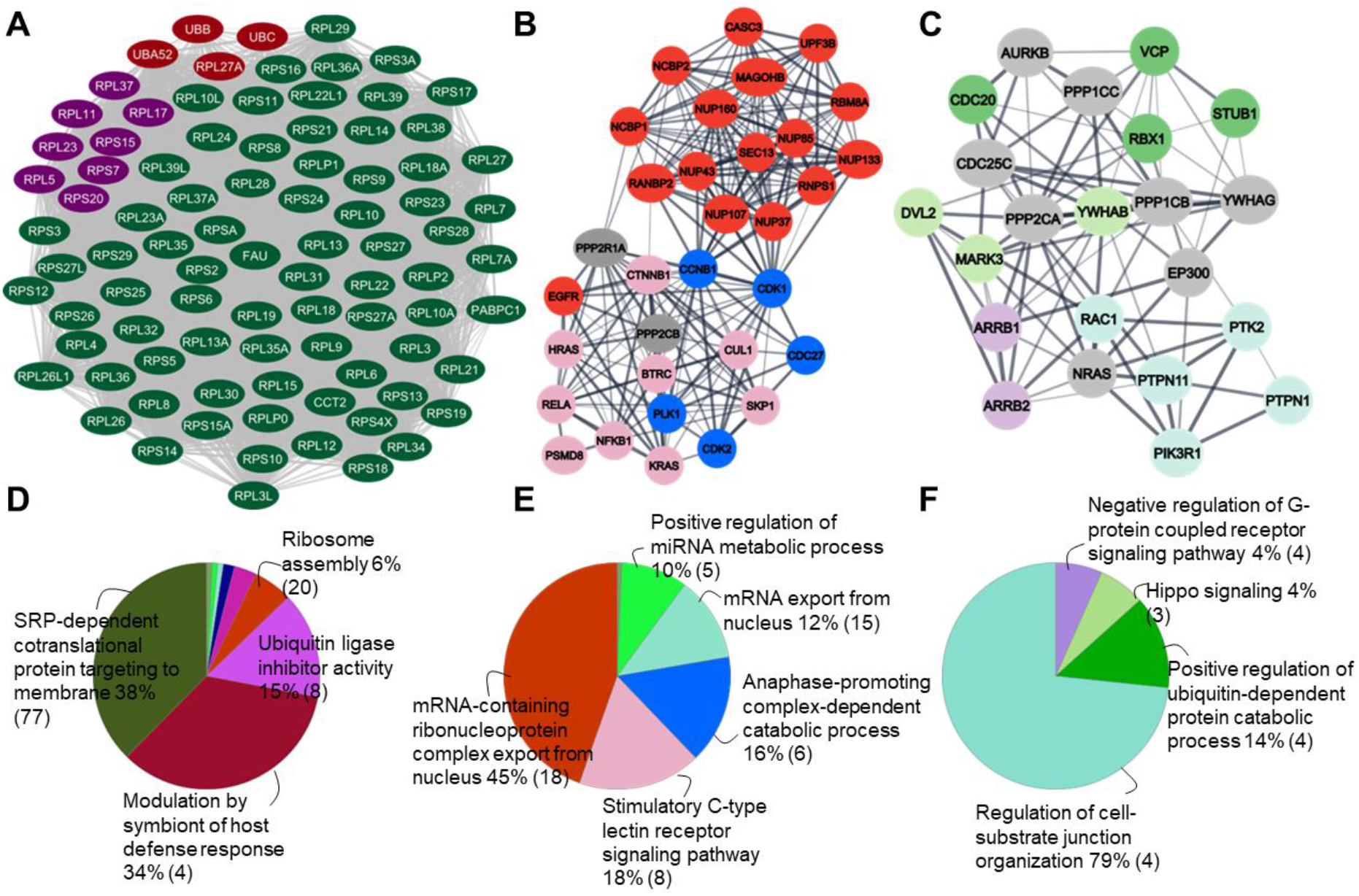
Function cluster of intermediate proteins. (A-C) PPI network of intermediate proteins that interact with protein phosphatases and viral protein generated in the MCode in three cluster accordingly with functional groups. (D-F) Especific agroupment in functional terms generate in the ClueGo. The number between parentheses are the genes number associated to that function (term). The percentual represent the most significant genes, the term with the most network leader the network interaction, represented by the functional groups. (A,D) Cluster 1; (B,E) Cluster 2; (C,F) Cluster 3.

In the second cluster, most proteins were shown to be involved in five signaling pathway: *i)* mRNA-containing ribonucleoprotein complex export from nucleus; *ii)* stimulatory C-type lectin receptor signaling pathway; *iii)* anaphase-promoting complex-dependent catabolic process; *iv)* mRNA export from nucleus; *v)* positive regulation of miRNA metabolic process (**Figure 4B**,**E**). While two of these functions are related to protein synthesis, one refers to cell mitosis.

In the third cluster, most proteins were shown to be involved in other four signaling pathway: *i)* regulation of cell-substrate junction organization; *ii)* positive regulation of ubiquitin-dependent protein catabolic process; *iii)* Hippo signaling; *iv)* negative regulation of G-protein coupled receptor signaling pathway (**Figure 4C,F**). In this cluster we also observed functions related to cell organization and protein degradation. Together those proteins work in cell organization, protein degradation and synthesis. It is known that the effect of SARS-CoV-2 infection promotes inflammatory response and cell damage. We suggest that the protein phosphatases together with intermediate proteins might be involved in the critical cellular processes to prevent cell damage caused by SARS-CoV-2 protein virus.

Taken together, our results suggest that protein serine threonine phosphatase 2A (PP2CA) and phosphatase tensin homolog (PTEN) seem to play an important biological role during SARS-CoV-2 infection, considering their interactions with various intermediate proteins (**Figure 3C,D, Supplementary Table 3**). We next proceeded for MCode and ClueGo analysis only with these PP2CA or PTEN network (**Figure 5**). For PP2CA, we observed two principal clusters, which the first cluster is the same for all proteins (**Figure 4A,D**) and the second involved in the mRNA export from nucleus and positive regulation of ubiquitin protein ligase activity (**Figure 5A**,**C**). For PTEN, we observed a principal cluster of interactions involved in the anaphase-promoting complex-dependent catabolic process, positive regulation of protein ubiquitination and phosphatidylinositol regulation (**Figure 5B,D**). We suggest the potential involvement of the PP2CA in protein turnover, and for PTEN in the mitosis and in the phosphatidylinositol signaling pathway.

**Figure 5:**
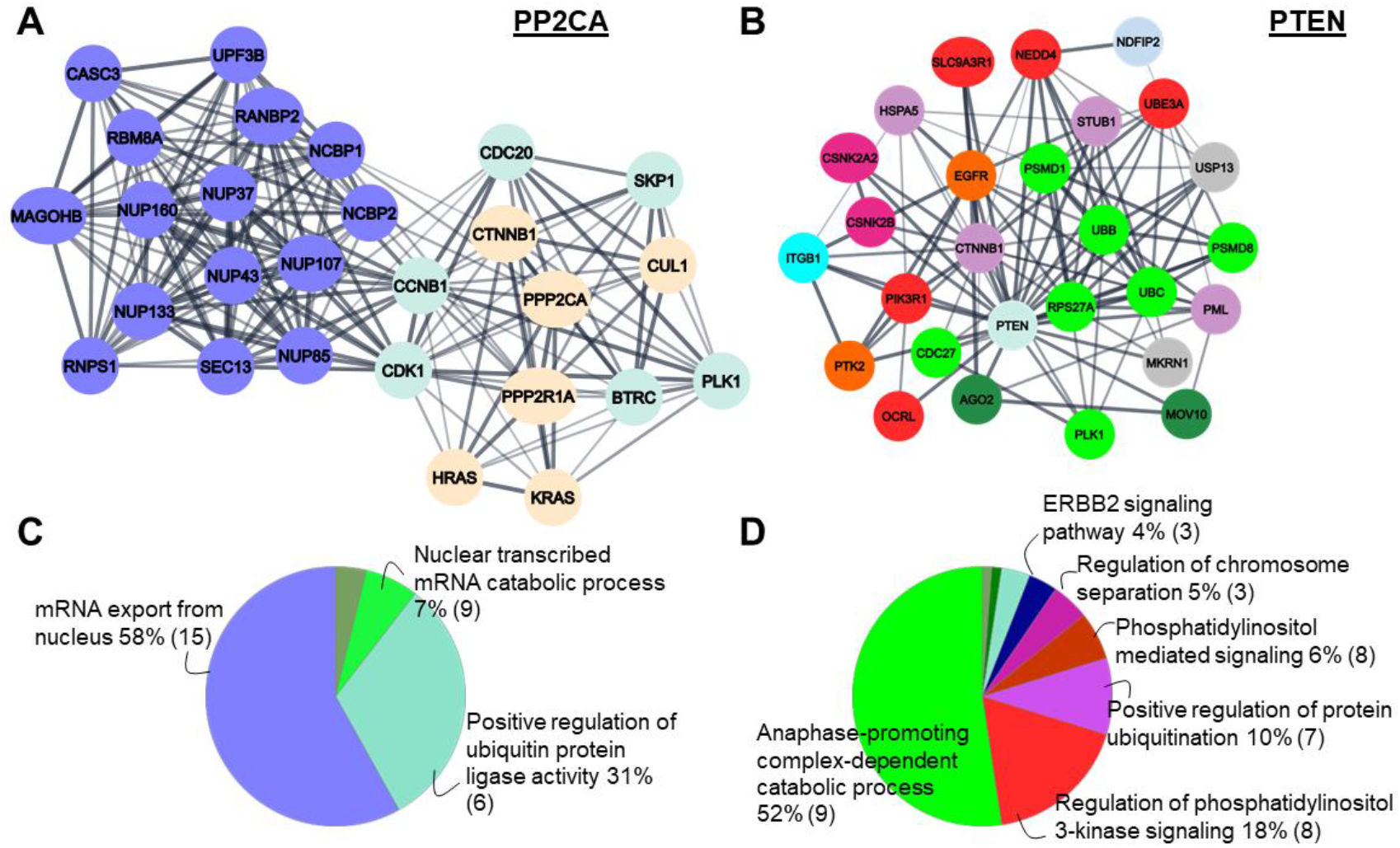
Function cluster of protein phosphatases PP2CA and PTEN. (A) PPI network of intermediate proteins that interact with protein phosphatase PP2CA and viral protein generated in the MCode in three cluster, accordingly with the functional groups, being the cluster 2 presented in the figure. (B) PPI network of intermediate proteins that interact with protein phosphatase PTEN generated in the Cytoscape. (C,D) Functional group generated in terms in the ClueGo. The number between parentheses are the genes number associated to that function (term). The percentual represent the most significant genes, the term with the leader of the network interaction, represented by the functional groups.

Export of messenger RNAs is known to be an important process for encoding antiviral factors in the body, and essential to produce proteins that are needed to counteract the virus replication and control infections ^42^. Thus, the enrichment of proteins involved in these function, may be related to the urgency of the body to fight the virus.

It is known that SARS-CoV-2 infection causes several processes of cell apoptosis as a result of the inflammatory response ^43^. One of the most obvious PTEN functions is to regulate mitosis, as well as proteins involved in the anaphase promoting complex-dependent catabolic process during infection. The anaphase promoting complex is a protein complex that mediate cycling degradation during anaphase, driving mitosis exiting and allowing cell growth. This complex also causes the degradation of proteins that hold the sister chromatids together, causing them to separate in anaphase and go to opposite poles of the cell ^44^. The existence of a large percentage of proteins involved in the anaphase promoting complex-dependent catabolic process during SARS-CoV-2 infection suggest that cell turnover is activated during SARS-CoV-2 infection as a mechanism to allow virus propagation.

In this sense, PTEN is also involved in the regulation of phosphatidylinositol 3-kinase signaling pathway. We observed that growth factor receptor signaling and PI3K was activated in Caco-2 cells and Huh7 cells infected by SARS-CoV-2, respectively ^7,12^. This activation results in Akt phosphorylation and subsequent activation of mTOR during SARS-CoV-2 infection in Huh7 cells ^7^, which regulate apoptosis, cell survival, host transcription and translation. Indeed, the viral replication decreases when the GFR signaling in inhibited thought pictilisib (PI3K inhibitor), omipalisib (PI3K and mTOR inhibitor) ^12^.

Therefore, understanding these cellular pathways will be important to the knowledge of COVID-19 infection mechanism and identification of different targets for the development of therapeutic agents for disease treatment.

### Phosphorylation level of intermediate proteins

Here we investigated the participation of protein phosphatases during viral infection. Then, we hypothesized that these proteins could act dephosphorylating the viral protein and/or the intermediate proteins involved in those signaling pathway described. To analyze the dephosphorylation of these proteins, we investigated the phosphorylation level of them in the phosphoproteome data performed in cells infected by SARS-CoV-2 ^6,11,19^. Once protein kinases phosphorylate and protein phosphatases dephosphorylate their substrates, we assume that the intermediate proteins with higher phosphorylation levels in the infected sample by SARS-CoV-2 compared to control have either higher protein kinase activity or lower protein phosphatase activity. Conversely, proteins with lower phosphorylation level in the infected sample by SARS-CoV-2 compared to the control indicate higher phosphatase activity and lower protein kinase activity.

To check if protein kinases are actually involved during SARS-CoV-2 infection, three studies analyzed the level of host phosphorylation proteins as well the kinases activities, extracted from vero E6 cells and A549 cells infected with the virus and nasopharinx swab sample from infected patient. These studies found 3036, 4643 and > 8500 human proteins/phosphorylation sites phosphorylated upon infection, which are involved in different cellular functions, such as cell cycle, apoptosis and DNA replication ^6,11,19^. They also identified activated and downregulated kinases upon infection and showed that kinases inhibition reduced viral replication ^6^. Interestingly, both studies identified phosphorylation sites in the SARS-CoV-2 proteins M, N, S, nsp3, nsp9, nsp14, and orf9b ^6,11^.

Analyzing the phosphorylation level of intermediate proteins identified, we observed lower phosphorylation level (protein phosphatase activity) in the FGFRTOP_S160 (1), RPS17_S113 (2), CDC20_T70 (3), ZNF318_S2091 (4), CENPF_S3054 (6), RANBP2_S2510 (7) and others in the model cells (**Figure 6A, Supplementary Table 4**). The intermediate proteins with higher phosphorylation level (protein kinase activity) observed in this study were MAPK1_Y187 and T185 (17, 18), RBM8A_S56 (19), MYCBP2_S3931, S3932 (20), RPS20_T9 (21) and others in the model cells (**Figure 6A, Supplementary Table 4**). Interestingly, in the patient sample we also observed lower phosphorylation level for RPS17_S113 (8), RANBP2_S2900 (1) and RPL and RPS family (3, 4, 5), while higher phosphorylation level for ZNF318 (12, 15, 17), but in different residues and PML proteins (14, 16) (**Figure 6B, Supplementary Table 4**). The protein RPS and RPL are involved in the SRP-dependent cotranslational protein targeting to membrane and ubiquitin ligase inhibitor activity function cluster as described above (**Figure 4A, D**) and mostly interacting in the network with PPP2CA (**Figure 3D**). Many works have been demonstrated phosphorylation/dephosphorylation regulation of RPS proteins, which acidic ribosomal (P1, P2 and P0) and RPS3 proteins could be dephosphorylated and regulated by PPP2CA, important in protein synthesis ^45,46^; RPS6 can be regulated by protein phosphatase 1, 2B (PPP1, PPP2B) activity after mitogenic stimulation ^47–49^; and RPL5 interact with PPP1 ^50^.

**Figure 6:**
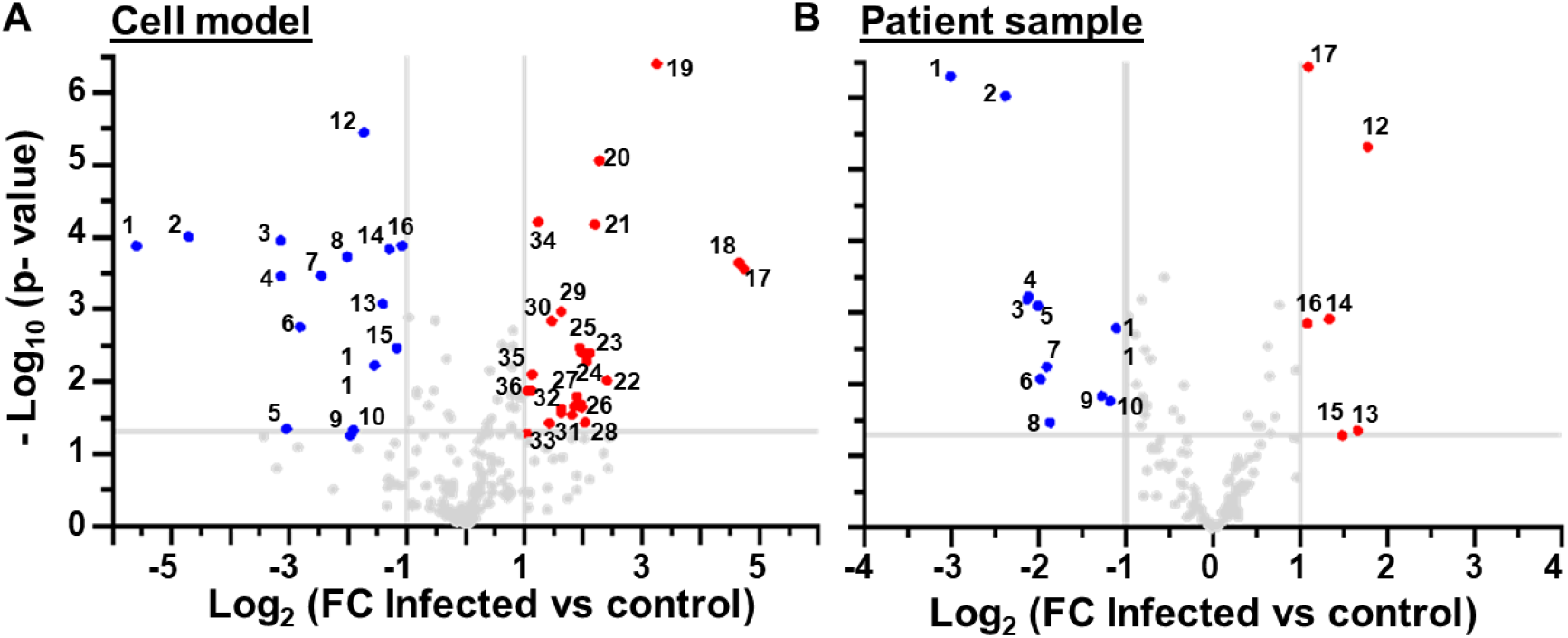
Phosphorylation degree of intermediate proteins. Volcano plot of phosphorylation level of intermediate proteins in (A) cell model and (B) patient sample. FC ≥ 1.0 represent the higher level of protein phosphorylation compared to control (red). FC ≤ 1.0 represent the lower level of protein phosphorylation compared to control (blue). FC = log2(infected/control). Graph was plotted using GraphPadPrism 9.0. Legend numbers in A, lower level of protein phosphorylation: 1- FGFRTOP_S160; 2- RPS17_S113; 3- CDC20_T70; 4- ZNF318_S2091; 5- UPF3B_S399; 6- CENPF_S3054; 7- RANBP2_S2510; 8- MARK3_S583; 9- GOLGA2_S37; 10- EP300_S1038; 11- RANBP2_T2153; 12- NDE1_S282; 13- RANBP2_S833, S837; 14- MARK3_S419; 15- CLASP1_S598; 16- SLC9A3R1_S290, S302 and higher level of protein phosphorylation: 17- MAPK1_Y187 ; 18- MAPK1_T185; 19- RBM8A_S56; 20- MYCBP2_S3931, S3932; 21- RPS20_T9; 22- PSMD1_T273 and PTK2_T273; 23- SLC9A3R1_S280; 24- SLC9A3R1_S269; 25- TOR1AIP1_S231 and ZNF318_S501; 26- MYCBP2_S1624; 27- CLASP1_S695 and RPL34_S12; 28- MPHOSPH10_S163, S167, S171; 29- RPS6_S235, S236; 30- EGFR_S991; 31- RPL30_S16 and RPL37_S96 and EGFR_S995; 32- MPHOSPH10_S167 and MPHOSPH10_S171 and RPS6_S244; 33- MYCBP2_S3478; 34- MYCBP2_S2687; 35- ZNF318_S2101; 36- RANBP2_T2458 and CSNK2B_S209. Legend numbers in B, lower level of protein phosphorylation: 1- RANBP2_S2900; 2- SKP1_T131; 3- RPLP1_S101; 4- RPLP2_S105; 5- RPSP1_S102; 6- VCP_S775; 7-NFKB1_S937; 8-RPS17_S113; 9- CTNNB1_S675; 10- ZNF318_S2101; 11- YWHAE_S210 and higher level of protein phosphorylation: 12- ZNF318_S305, S590, S250, S134, S307, S592, S252; 13- RPS6_S240; 14- PML_S480; 15- ZNF318_S69; 16- PML_S504; 17- ZNF318_S4.

The intermediate proteins with lower phosphorylation level seems to be dephosphorylated specially by PPP2CA (**Supplementary Figure 1**) once we observed interaction between them (**Figure 3D**). While the intermediate proteins with higher phosphorylation levels should be phosphorylated by protein kinases, with lower participation of the protein phosphatases PPP2CA, PTEN, PPP1CB, PPP1CC and PTPN12 (**Supplementary Figure 1**).

SARS-CoV-2 is known to cause changes in the host cells that can lead to apoptosis, making the organism to produce new cells and use the defense system to fight the virus ^43^. CDC20 is involved in mitosis checkpoint, acting as a regulator of cell division through ubiquitination ^51,52^. If the protein phosphatase is involved in the virus infection in the sense that it interacts with the virus and other intermediate proteins, it may interfere with the activity of CDC20 in a way that induces new cell formation. Since CDC20 is involved in the function of regulating cell division. Its known that the protein phosphatase PPP2CA is essential for life cycle of virus due its importance in controlling the cell cycle and apoptosis^53^. Additionally, PPP2A is important for the activation of the anaphase promoting complex (APC/C), suggesting that dephosphorylation of CDC20 by PPP2CA is required for the complex to be activated ^54–56^. In terms of infection, if CDC20 has decreased phosphorylation during infection, it is inferred that APC/C cannot be activated, or will be activated incorrectly, leading to cell cycle arrest and/or cell apoptosis. In addition, SARS-CoV-2 infection promotes activation of p38/MAPK and Casein kinase 2 (CK2) signaling activity and shut down mitotic kinases, resulting in cell cycle arrest ^6^.

These studies indicated the balance between protein kinases and phosphatases during cell infection and viral replication. Therefore, more studies become necessary to demonstrate the direct role of protein kinase and phosphatase regulating SARS-CoV-2 infection and replication in human cells.

## Conclusions

During SARS-CoV-2 infections, host and virus proteins interactions are essential for the onset of viral replication. In this work we demonstrated the changing in the protein phosphatase levels, which protein tyrosine phosphatase and protein serine/threonine phosphatase families seems to be higher and lower expressed, respectively during SARS-CoV-2 infection. We identified protein interactions between protein phosphatases and SARS-CoV-2 proteins. By expanding the database searches, we also found numerous intermediate proteins participating in the interaction network. This suggest the existence of an important relationship between these proteins, since each one of them participates in a specific signaling pathway. These interactions may also interfere in the immune response in different ways, and potentially participate in the different observed symptoms.

We identified two protein phosphatases with more evident interactions, PPP2CA and PTEN. Among all the functions, the most evident were the export of mRNA to the nucleus, and anaphase promoter complex-dependent catabolic process. This, led us to infer that while the virus infects our cells and causes inflammatory and apoptotic processes, the cell turnover is activated in an attempt to prevent excessive loss of cells and tissue damage.

We also identified the intermediate proteins RBM8A, RPS proteins and CDC20 with very significant variations in their phosphorylation levels, which seems to be the most affected by infection. Interestingly, RPS proteins have been demonstrated to be regulated by protein serine threonine phosphatases and CDC20 interacts with the phosphatase PPP2CA. This regulation and interaction are necessary for the protein synthesis and anaphase promoter complex to be activated, respectively. Based on the importance of PPP2CA to be modulated by different virus, the data reported here suggest the modulation of this protein also by SARS-CoV-2 virus.

Together with the protein kinases, which phosphorylate substrates, the protein phosphatases are important for dephosphorylating specific sites in this complex, regulating cell signaling. These proteins and all the others identified in the study may undergo changes in their signaling pathways caused by modifications in their phosphorylation levels, due the expression level and phosphatase activity of protein phosphatases.

## Acknowledgements

We would like to thank Dr. Marcus Oliveira and Dr. Francisco Prosdocimi for the critical reading of this manuscript. This work was supported by grant 2019/02605-3, the fellowship 2021/10474-6 and 2020/10168-0 from São Paulo Research Foundation (FAPESP) and PIBIC 2021/2022 from National council for Scientific and Technological Development (CNPq).

## Conflict of interest

The authors declare no conflict of interest.

## Author contributions

G.I.M.G., L.P.O. and L.E.S.F.M. contributed to data analysis and to figure drawing and editing. All authors conceptualized the study, wrote and prepared the original draft, critically reviewed and edit the manuscript. All authors reviewed the results and approved the final version of the manuscript.

## Notes

### Competing Interest Statement

The authors have declared no competing interest.

