## Supplementary Figure 1 for "Potential involvement of protein phosphatase PP2CA on protein synthesis and cell cycle during SARS-CoV-2 infection. A meta analysis investigation"

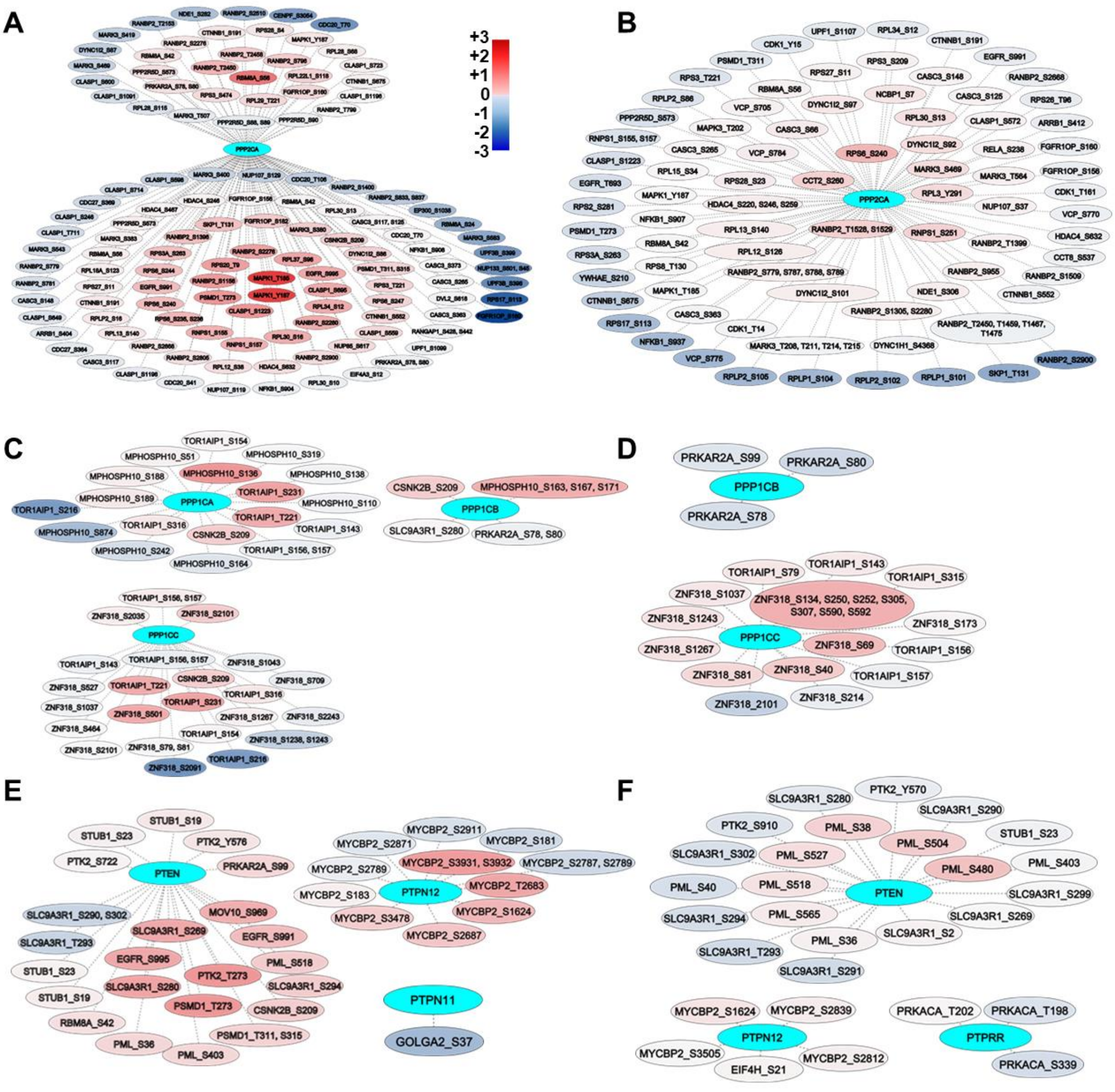


**Supplemmentary Figure 1: Phosphorylation degree of intermediate proteins network.** Phosphorylation level by color of intermediate protein that interact with protein phosphatase, made in Cytoscape software in (A,C,E) cell model and (B,D,F) patient samples. The proteins with higher and lower level of phosphorylation are in red and blue, respectively, accordingly with the phosphorylation degree. The color scale is in the figure.
